# Sensitive, highly multiplexed sequencing of microhaplotypes from the *Plasmodium falciparum* heterozygome

**DOI:** 10.1101/2020.02.25.964536

**Authors:** Sofonias K Tessema, Nicholas J Hathaway, Noam B Teyssier, Maxwell Murphy, Anna Chen, Ozkan Aydemir, Elias M Duarte, Wilson Simone, James Colborn, Francisco Saute, Emily Crawford, Pedro Aide, Jeffrey A Bailey, Bryan Greenhouse

**Author notes:** The first two authors contributed equally to this work. The last two authors contributed equally to this work.

## Abstract

**Background:** Targeted next generation sequencing offers the potential for consistent, deep coverage of information rich genomic regions to characterize polyclonal *Plasmodium falciparum* infections. However, methods to identify and sequence these genomic regions are currently limited.

**Methods:** A bioinformatic pipeline and multiplex methods were developed to identify and simultaneously sequence 100 targets and applied to dried blood spot (DBS) controls and field isolates from Mozambique. For comparison, WGS data were generated for the same controls.

**Results:** Using publicly available genomes, 4465 high diversity genomic regions suited for targeted sequencing were identified, representing the *P. falciparum* heterozygome. For this study, 93 microhaplotypes with high diversity (median H_E_ = 0.7) were selected along with 7 drug resistance loci. The sequencing method achieved very high coverage (median 99%), specificity (99.8%) and sensitivity (90% for haplotypes with 5% within sample frequency in DBS with 100 parasites/µL). In silico analyses revealed that microhaplotypes provided much higher resolution to discriminate related from unrelated polyclonal infections than biallelic SNP barcodes.

**Discussion:** The bioinformatic and laboratory methods outlined here provide a flexible tool for efficient, low-cost, high throughput interrogation of the *P. falciparum* genome, and can be tailored to simultaneously address multiple questions of interest in various epidemiological settings.

## INTRODUCTION

Malaria genomics have been applied to generate actionable data to inform control and elimination efforts, e.g. tracking the spread of drug resistance and evaluating the response to vaccine candidates [1–3]. To realize the full potential of genomics for understanding malaria transmission, an ideal genotyping method would seek to maximize discrimination between infections, including polyclonal, low density infections often encountered in endemic settings. Traditional genotyping methods such as typing of length polymorphisms [4], microsatellites [5,6] and single nucleotide polymorphisms (SNPs) [7–9] have been extensively used to characterize malaria transmission. However, technical and biological constraints limit the scalability and discriminatory resolution of these methods, particularly when infections are polyclonal [10,11]. Typing of microsatellites and length polymorphisms suffer from difficulties in standardizing laboratory protocols, allele calling and reporting, and detection of minority clones [12]. To address challenges in throughput and standardization, several SNP barcoding approaches were developed to evaluate parasite diversity and population structure and to estimate transmission dynamics [7–9,11,13]. However, SNP-based methods have limited discriminatory power to compare polyclonal infections, which represent the majority of the parasite population in many places in sub-Saharan Africa, including areas of low transmission [14,15].

Recent advances in next-generation sequencing (NGS) allow targeted deep sequencing of short, highly variable regions with numerous alleles (microhaplotypes) [16,17], predominantly composed of 3 or more SNPs, allowing for detailed characterization of the ensemble of parasites in an infection. Most applications of these methods to date have targeted one or a few genomic loci to provide information on drug resistance, composition of infections or selection [18–25]. Extending these methods to numerous genetically diverse loci offers the potential for high resolution comparisons of infections at a population level [17]. To this end, there have recently been efforts to multiplex large numbers of loci that reflect overall patterns of *P. falciparum* diversity [26,27].

With thousands of whole-genome sequencing (WGS) data available [28,29], it is now possible to establish an optimal set of multi-allelic targets to interrogate the *P. falciparum* genome using NGS. However, bioinformatics pipelines to identify informative targets are currently lacking. Furthermore, sensitive and high throughput laboratory methods for targeted sequencing of these markers in a cost-effective manner remains a major challenge. In this study, we describe the initial evaluation of a bioinformatic pipeline to identify high value *P. falciparum* microhaplotypes and a robust PCR-based laboratory method that allows sequencing of hundreds of these microhaplotypes in a single reaction. This study evaluates the performance of the laboratory method and the information content of a selection of ∼100 microhaplotypes across a range of in silico analyses, mixture controls, and field samples.

## METHODS

### Microhaplotype selection pipeline

Tandem repeats longer than 50bp within the *P. falciparum* 3D7 reference genome were determined using tandem repeat finder [30]. After excluding tandem repeats, overlapping windows of 200bp every 100bp were created. Per base coverage and the fraction of proper pairs (i.e. the number of pairs with both mates mapped with normal insert sizes in the proper orientation divided by the total number of pairs mapped to a region) were calculated from the raw Illumina data of 13 reference genomes [31]. Windows were kept if they were within 1 standard deviation from the average base-pair coverage of the genome and if the proper pair fraction was greater than 0.85 in 11 out of the 13 genomes. Windows with dinucleotide repeats or homopolymers longer than 10bp were excluded. The 3D7 sequences from these windows were then blasted against the 13 genomes with at least 75% identity and 95% coverage using LASTZ [32]. Windows were kept if they mapped once to all 13 genomes, had no length variation greater than 3 bases, average GC content >15%, and sequences in at least 3 of the 13 genomes were unique. Local haplotype assembly was run on these final windows in WGS from 4054 field samples (Supplementary Table 1) and 33 lab isolates).

### Mock and field DBS samples

Mock DBS samples were prepared as previously described [33]. Briefly, synchronized *P. falciparum* parasites were mixed at different proportions with uninfected human whole blood to obtain a range of parasite densities (10, 100, 1000 and 10000 parasites per μL of blood) (Supplementary Figure 1). Field DBS samples were collected from southern Mozambique from febrile malaria cases. This study was approved by the Institutional Review Board at the University of California San Francisco and Manhiça Health Research Centre (CISM), Mozambique. DBS samples were stored at −20°C until processing. DNA was extracted from a single 6mm hole-punch using a modified Tween-Chelex protocol as previously described [33].

### Multiplexed amplicon sequencing

Primers were designed for 100 selected genomic regions of the microhaplotypes and drug resistance markers using the CleanPlex^®^ algorithm (Paragon Genomics Inc, USA). Amplification of the 200-plex oligo pool was performed with some modifications of the CleanPlex® protocol (Paragon Genomics Inc, USA) (see Supplementary Methods). Samples were bead purified, quantified and pooled. The final library was bead purified, assessed for quality and sequenced with 150bp paired end clusters on a NextSeq instrument (Illumina, San Diego, CA, USA). The targeted amplicon data were analyzed using SeekDeep (v.2.6.6) [34] (see Supplementary Methods).

### Selective whole genome amplification and whole-genome sequencing

Selective whole genome amplification and whole-genome sequencing were performed following a previously optimized protocol [33]. After demultiplexing, high quality reads were aligned to the *P. falciparum* 3D7 reference genome (version 3) with BWA-MEM [35]. Variants were called using the Genome Analysis Toolkit (GATK) Best Practices [36].

### Simulations to evaluate genetic relatedness

Population level allele frequencies were calculated for 91 microhaplotypes and variant loci across Pf3k samples from Ghana, due to the large number of publicly available WGS in Africa. Simulated genomes were created with a single multinomial draw given allele frequencies across all loci. Relatedness between genomes was simulated parameterizing a Bernoulli draw by the expected relatedness over the number of loci, masking the selected loci, and making both genome pairs equivalent at each masked locus. Relatedness was calculated by converting each locus to a boolean vector, concatenating all loci in a sample to a single vector, then calculating Jaccard distance between sample pairs.

## RESULTS

### The heterozygome of P. falciparum: identification of high diversity genomic regions

A custom pipeline was developed to perform a genome-wide scan of *P. falciparum* variation and identify short genomic regions that exhibit high population level genetic diversity, here termed ‘heterozygome’ windows (Figure 1A). As a first stage, we calculated overlapping windows of 200bp excluding regions with tandem repeats. Windows that contained no homopolymers, dinucleotide repeats longer than 10bp, or length variation greater than 3bp were considered to be suitable for downstream PCR and sequencing. Windows were kept if they uniquely mapped to each of the 13 high-quality assembled and annotated genomes of *P. falciparum* isolates [31] with at least 70% identity. Out of these 63,414 windows, 4,465 windows had at least 3 unique haplotypes in 13 reference genomes and were retained as part of the heterozygome. Further population genetic measures were calculated for the heterozygome windows by running local haplotype reconstruction for each window on 4054 field samples with publicly available WGS [28,37–41], broadinstitute.org] (Supplementary Table 1).

**Figure 1.**
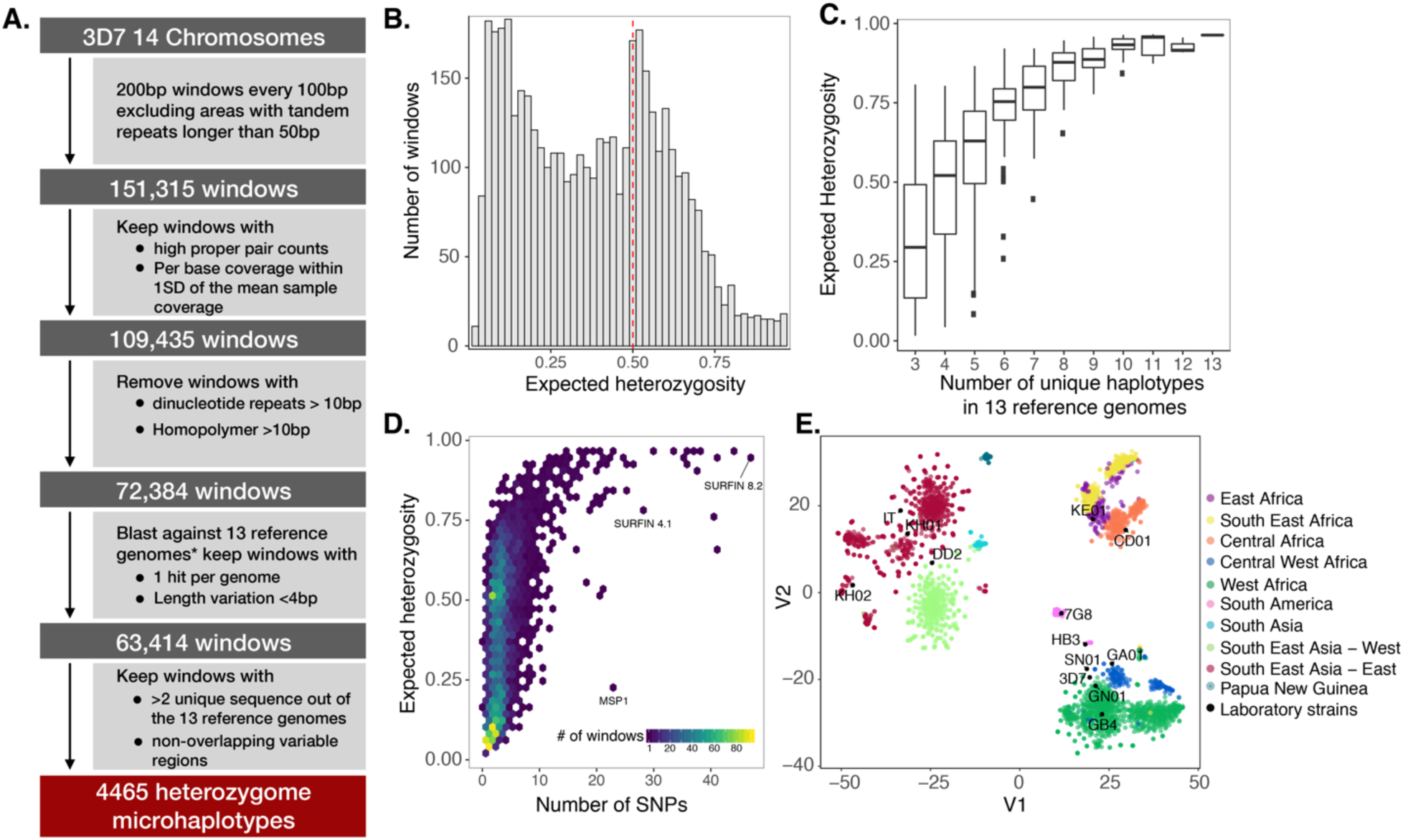
Characterization of the heterozygome of *P. falciparum* in global parasite populations. **A.** Bioinformatics workflow for the identification of the heterozygome. *13 high-quality assembled and annotated genomes of *P. falciparum* isolates [31]. The variable region covered was determined as the first position to the last position within the window with a SNP or INDEL >=0.5%. The window with the highest number of samples was kept when variable regions overlapped. **B.** Distribution of expected heterozygosity of microhaplotypes (n=4465) in global parasite populations (n=4054 isolates), showing high genetic diversity in a substantial number of microhaplotypes. **C.** Relationship between the number of unique haplotypes in 13 reference genomes and the expected heterozygosity of microhaplotypes in global parasite populations. **D.** Distribution of the number of SNPs per microhaplotype, showing the relationship between the number of SNPs and expected heterozygosity. **E.** Population structure of the global *P. falciparum* parasite population inferred from the microhaplotypes visualized using tSNE as implemented in the Rtsne package [42] with 25,000 iterations, a perplexity parameter of 100, and a trade-off θ of 0.5.

A third of the microhaplotypes in the heterozygome had high expected heterozygosity (H_E_ > 0.5, 35%) in the global parasite population, and 356 had H_E_ > 0.7 (Figure 1B). Heterozygosity was correlated with the number of unique haplotypes in the 13 reference genomes used (Spearman rho = 0.5, p-value < 0.001, Figure 1C), confirming the utility of this initial filtering approach. Geographically, the highest diversity was observed in West and Central West Africa (Supplementary Table 2). More than half (55%) of the microhaplotypes were composed of at least 3 SNPs (Figure 1D) and 846 microhaplotypes had at least 5 SNPs. Heterozygome microhaplotypes were able to accurately represent geographic structure of the global parasite population (Figure 1E).

### Selected microhaplotypes for multiplexed sequencing

The 4,465 microhaplotypes were ranked by within-Africa expected heterozygosity and genetic differentiation (Jost’s D). The 150 most diverse and differentiated microhaplotype loci with identifiable flanking primer sites were combined with selected molecular markers of drug resistance (n = 11) for primer design. For this study, 100 loci (93 microhaplotypes and 7 markers of drug resistance) were selected and multiplexed (Figure 2A). The selected microhaplotypes were genetically diverse in all malaria-endemic regions, with a median heterozygosity ranging from 0.48 in South America to 0.67 in Central Africa. On average, there were 5 (IQR: 3 - 7) SNPs and 3.4 (IQR 2.8 - 4.0) effective alleles in the selected microhaplotypes (Figure 2B). A subset of the microhaplotype loci were highly differentiated between malaria endemic regions and between countries within a region (Figure 2C), leading to strong population structure (Figure 2D). The selected microhaplotype loci were distributed throughout the 14 chromosomes of the parasite (median = 6, range 2-11 loci per chromosome) (Figure 2E, Supplementary Table 3).

**Figure 2.**
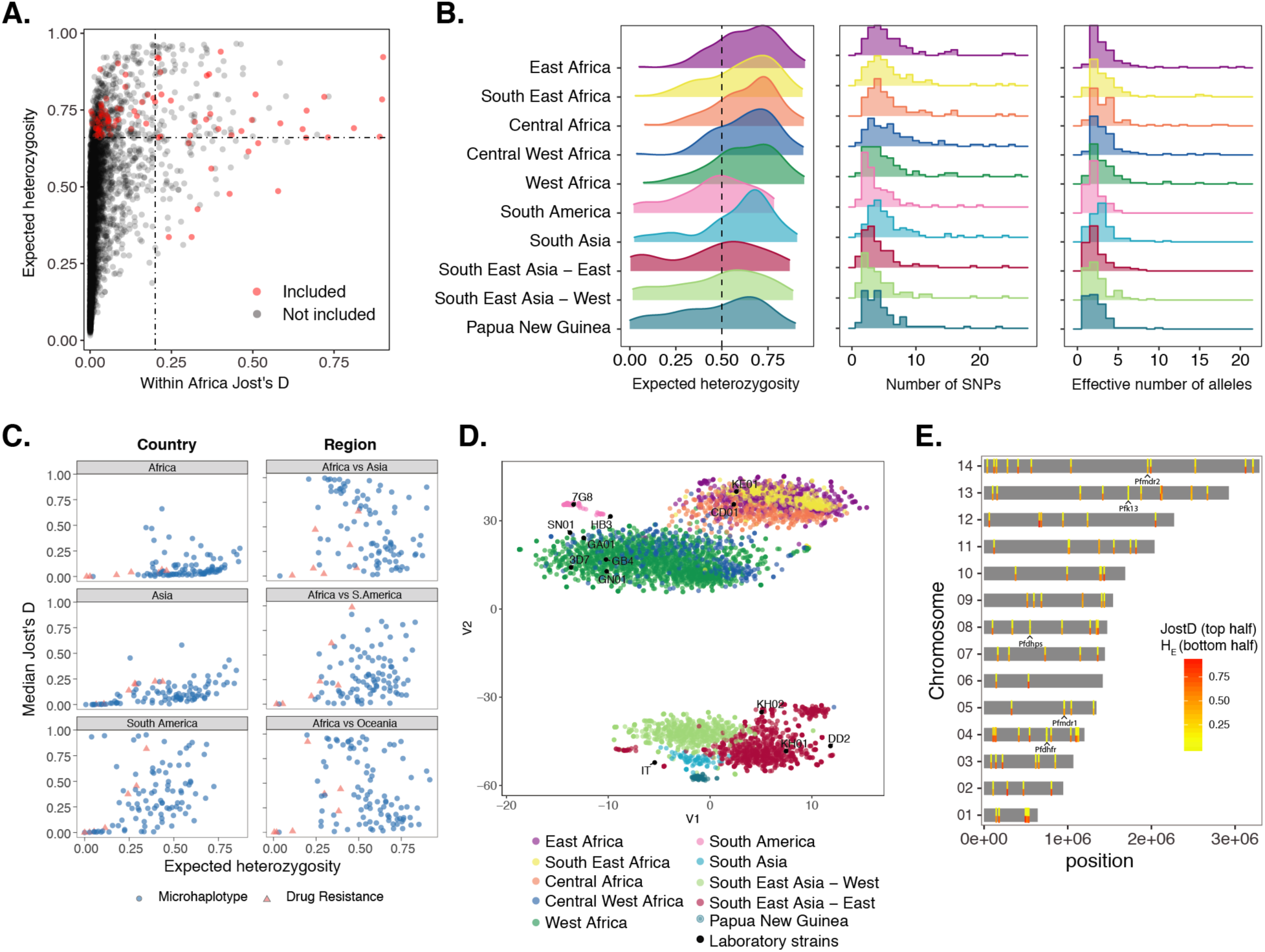
Overview of identified and selected microhaplotype loci (n=93) and drug resistance markers (n=7). **A.** Relationship between expected heterozygosity and within-Africa genetic differentiation (Jost’s D) for selected microhaplotypes. Microhaplotype loci were screened for identifiable primer sites before primer design. **B.** Distribution of expected heterozygosity, number of SNPs and the number of effective alleles of the selected microhaplotype loci in different malaria endemic regions. **C.** Relationship between expected heterozygosity and genetic differentiation between countries within regions and between different regions. **D.** Geographic clustering of global parasite populations inferred from the selected microhaplotypes visualized using tSNE as implemented in the Rtsne package [42] with 25,000 iterations, a perplexity parameter of 100, and a trade-off θ of 0.5. **E.** Chromosomal location of microhaplotypes. The mean of within-Africa genetic differentiation (top half) and mean expected heterozygosity (bottom half) are shown by coloured bars. The molecular markers of drug resistance included in this panel are also indicated.

### Multiplexed targeted sequencing of microhaplotype and drug resistance markers

A two-step multiplex PCR based assay was optimized using CleanPlex^®^ chemistry (Paragon Genomics Inc, USA) for evenness of coverage and detection of minority clones. The assay was evaluated on a range of dried blood spot controls containing known proportions of laboratory strains (Supplementary Figure 1). A high level of uniformity in coverage was achieved across different parasite densities (Figure 3A), with detection of alleles ranging from 93 to 100% in samples down to 10 parasites/µL (Figure 3B). The assay demonstrated high specificity regardless of the proportions of strains in the mixture (true positive rate = 99.8%). The sensitivity of the method was high across a wide range of parasite densities (Figure 3C). For example, at a within-sample haplotype proportion of 0.05, an average sensitivity of 61% was observed for 10 parasites/µL total parasite density and 90% for 100 parasites/µL, indicating the suitability of the method for the detection of minority clones even in low density infections (Figure 3C). Finally, the high correlation between the observed and expected haplotype proportions indicates the potential of this method for accurate quantification of within-host proportions of strains in mixed infections (Figure 3D).

**Figure 3.**
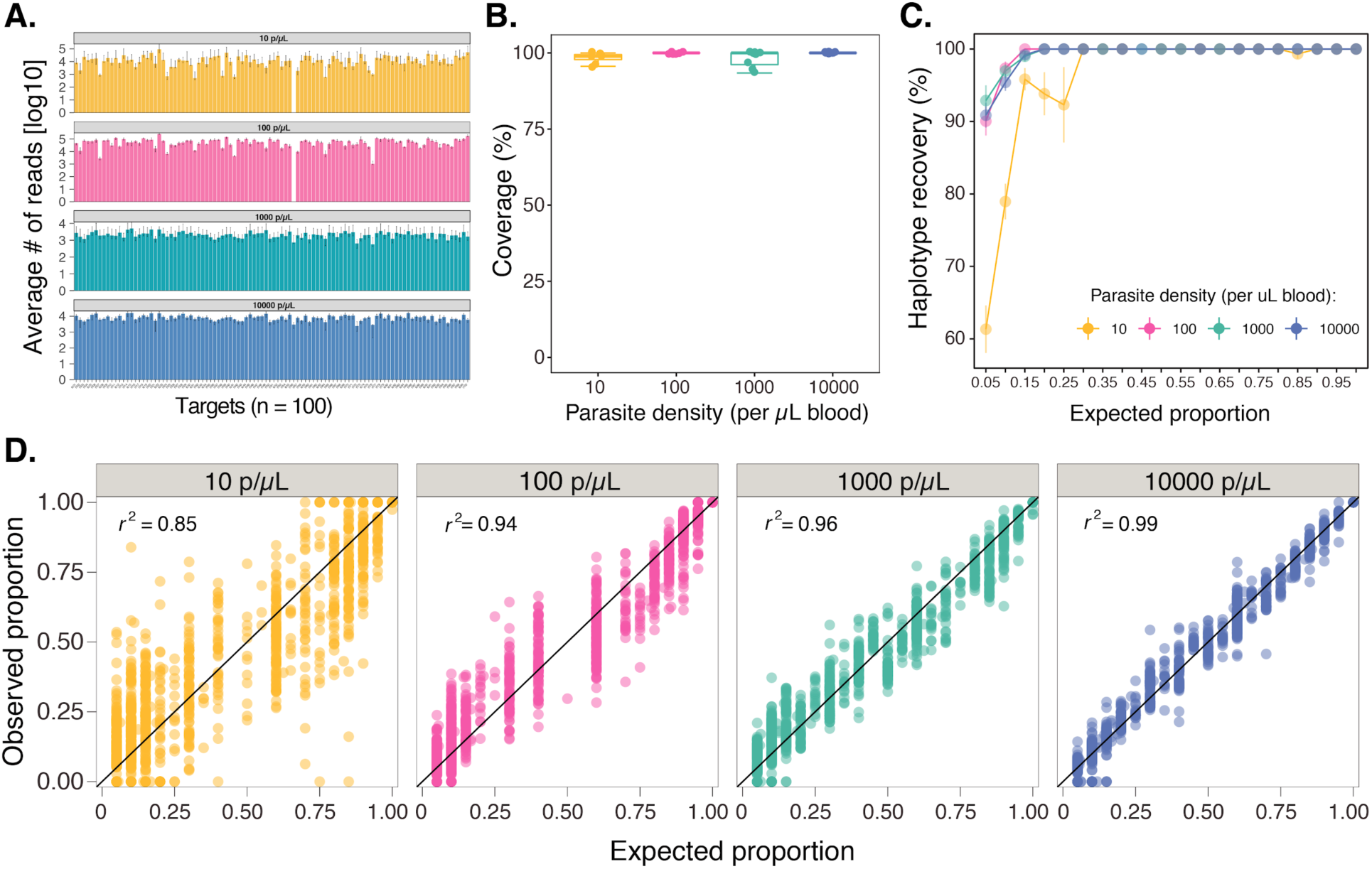
Coverage and sensitivity of multiplexed targeted sequencing of microhaplotypes (n=93) and drug resistance markers (n=7) on control samples. **A.** Average number of reads per target per sample. The median (bars) and interquartile range (error bars) are shown. **B.** Boxplot summarising the coverage of microhaplotype loci and drug resistance targets by parasite density. Coverage was determined based on the number of targets with 250 or more reads. **C.** Sensitivity of the assay for the detection of haplotypes at different proportions stratified by total sample parasite density is shown. Microhaplotype recovery was calculated as the number of observed haplotypes matching to the expected microhaplotypes divided by the total number of expected haplotypes. **D.** Correlation of expected and observed within-sample proportions of microhaplotypes by parasite density is shown.

### Targeted sequencing provides substantially better detection of microhaplotypes than whole genome sequencing

Compared with targeted sequencing, whole genome sequencing (WGS) provides broader evaluation of the genome but at the expense of sequence depth in specific regions of interest. To compare the sensitivity of targeted vs. WGS for detection of microhaplotype loci, WGS data were generated from the same DBS mixture controls, using selective whole genome amplification, at high depth and coverage (Supplementary figure 2). From these WGS data, reads spanning the entire variable regions of microhaplotypes were extracted to obtain unambiguous sequences. The extracted data were analyzed to evaluate coverage, detection of minority clones and quantification of strains. In contrast to results from targeted sequencing, microhaplotypes extracted from WGS data had poor coverage and sensitivity for the detection of minority alleles, even at the highest parasite densities evaluated (Supplementary figure 3).

### Microhaplotypes provide accurate estimation of complexity of infection (COI) and genetic relatedness

To compare the accuracy of COI estimation and genetic relatedness between infections by microhaplotype and SNP-based approaches, we extracted molecular SNP barcodes (n=24, [7]), Spot Malaria SNPs (n=87 of the 101 SpotMalaria SNPs [9,29]) and 100 high coverage and high diversity SNPs with > 0.2 minor allele frequency from the WGS data generated on the controls. Compared to biallelic SNPs, targeted sequencing of microhaplotypes consistently achieved substantially more accurate estimation of COI and, even more dramatically, pairwise genetic relatedness at all parasite densities, indicating the high discriminatory power of microhaplotypes for the measurement of within-host diversity and genetic relatedness between infections (Figure 4). These findings are consistent with the high genetic diversity of the selected microhaplotype loci and the high sensitivity of the assay for detecting minority clones.

**Figure 4.**
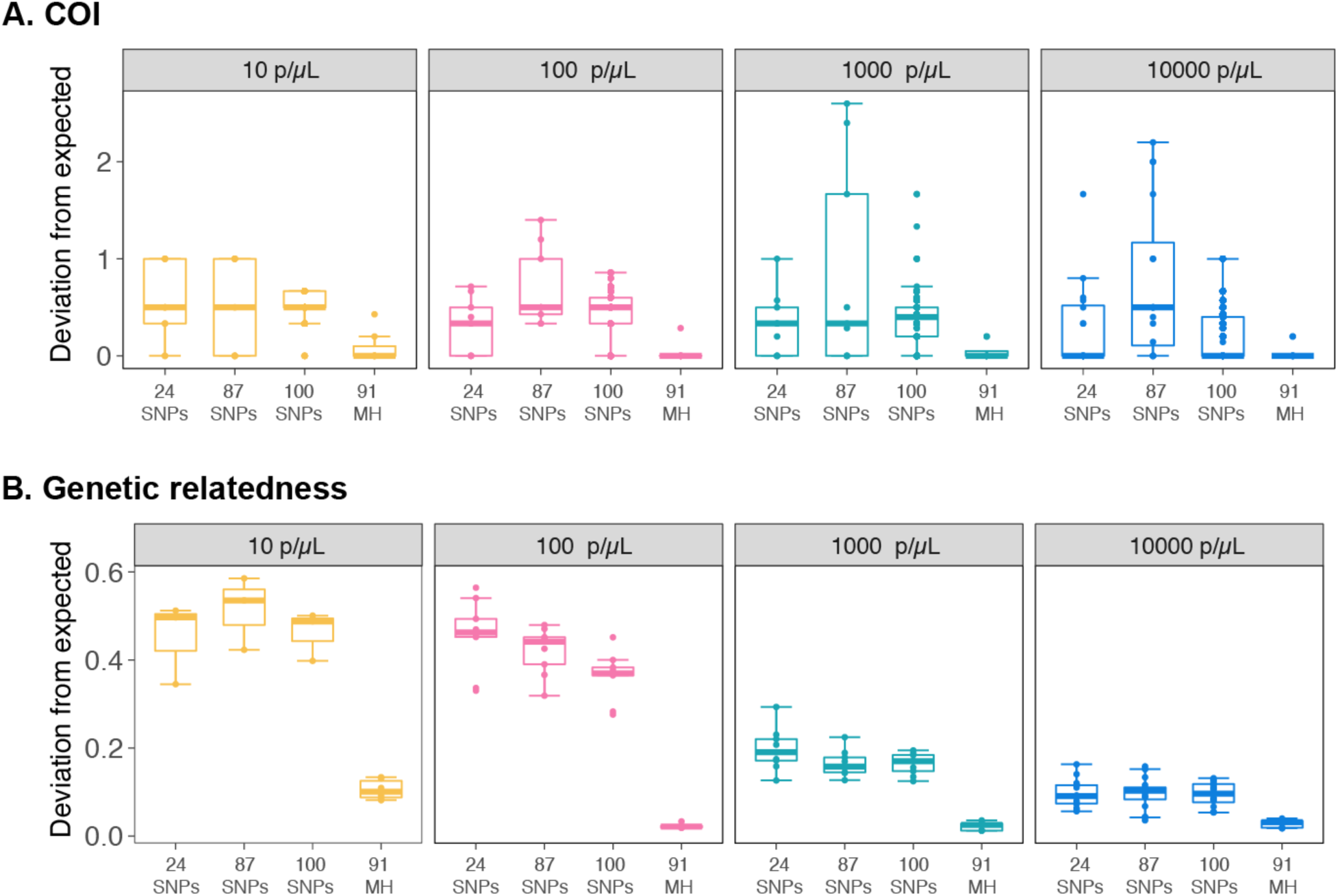
Comparisons of marker sets for complexity of infection (COI) and pairwise genetic relatedness. **A.** The deviation of observed COI from the expected number of strains in the mixture controls. COI was determined using THE REAL McCOIL [43] for SNPs and the number of unique haplotypes per samples for microhaplotypes. **B.** The deviation of observed genetic relatedness from the expected. The observed pairwise genetic relatedness was determined using Jaccard distance and is compared with the expected genetic relatedness between the control samples. Deviation was calculated as mean absolute error. 24 SNP barcodes, 87 SNPs and 100 high coverage and high diversity SNPs were extracted from the WGS data. 91 high quality microhaplotype loci (MH) were used for these analyses.

### Microhaplotypes provide higher discriminatory power than individual SNPs for estimating genetic relatedness of polyclonal infections

Polyclonal infections are common in many endemic areas, making it difficult to estimate genetic relatedness between infections. In order to evaluate the discriminatory power of microhaplotypes vs. SNPs, a simple simulation of infection pairs with varying numbers of clones and related parasites was performed. Perfect detection of alleles was assumed, to isolate the information content of the loci from the sensitivity of the laboratory method. For monoclonal infections, all methods performed similarly and were able to easily discriminate unrelated parasites from siblings and clones (Figure 5). However, in polyclonal infections, microhaplotypes provided higher genetic resolution to discriminate related from unrelated infections across all scenarios evaluated. We note that that the simulation framework and distance metric used here were intentionally straightforward in order to convey a high-level comparison of the information content of the various sets of loci, and do not represent a comprehensive, quantitative evaluation using methods directly tailored at inferring ancestry.

**Figure 5.**
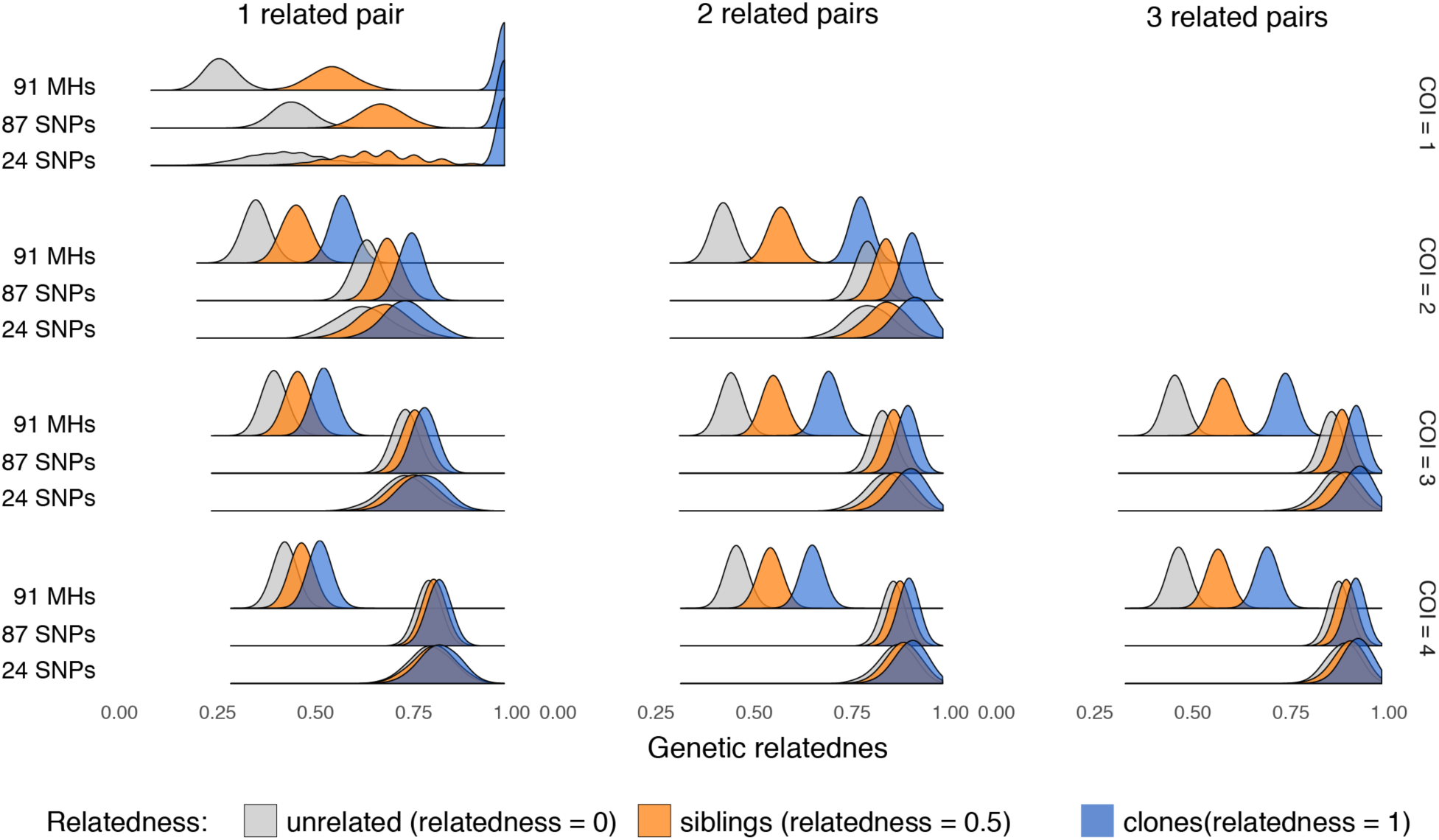
In silico comparison of genetic relatedness estimated from multi-allelic and biallelic loci in polyclonal infections. Peaks represent distributions of genetic distance obtained from 2000 simulated infection pairs for each combination of COI, relatedness, and, set of loci. Separation between the peaks demonstrates discriminatory power among unrelated, sibling, and clones of parasites using 91 microhaplotypes (91 MHs), 87 Spot Malaria SNPs, and 24 SNP barcodes.

### Validation of multiplexed amplicon sequencing in field samples

82 *P. falciparum* DBS samples from southern Mozambique were genotyped using the microhaplotype panel. Similar to the control samples, a high level of uniformity in the average number of reads per target (Figure 6A) and a coverage of > 95% was achieved in samples with at least 10 parasites/µL (Figure 6B). Analyses of replicate DBS samples extracted and processed independently showed the reproducibility of the assay for the detection and quantification of haplotypes in monoclonal and polyclonal samples (Figure 6C). The majority (68%) of the infections were polyclonal with a mean COI of 2.3 (Figure 6D). The selected microhaplotypes were also diverse in the three provinces of southern Mozambique (average heterozygosity = 0.6 and average effective alleles = 2.6), and 72 of the 93 microhaplotypes had H_E_ > 0.5 in at least one of the three provinces (Figure 6E). There was strong correlation (r^2^ = 0.74, p < 0.001) between the heterozygosity of the microhaplotypes in the global parasite population and the observed heterozygosity in Mozambique (Figure 6F), confirming the validity of the pipeline to identify high diversity genomic regions even in those countries that did not have publicly available WGS for the target selection. Visualization of microhaplotypes and biallelic SNPs (obtained from WGS data) on the same polyclonal samples illustrated the potential of microhaplotypes to better characterize these infections, given the large number of heterozygous SNP calls (Figure 6G). Analyses of drug resistance targets that were multiplexed in this assay showed absence of known resistance-associated K13 mutations in this population.

**Figure 6.**
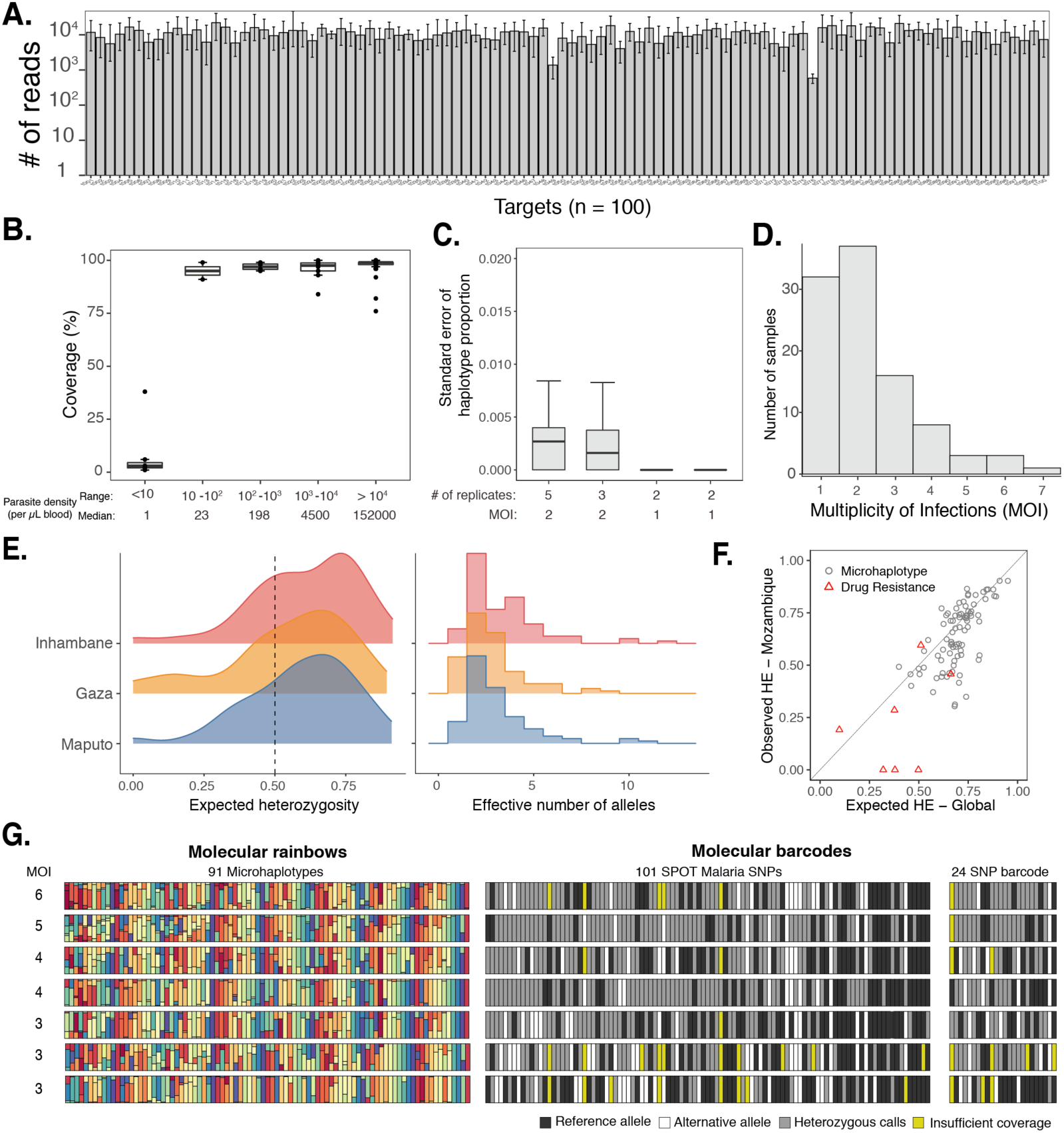
Coverage of multiplexing assay on selected microhaplotypes (n=93) and drug resistance markers (n=7) on field samples. **A.** Average number of reads per target per sample. The median (bars) and interquartile range (error bars) are shown for 82 field samples. **B.** Boxplots summarize the coverage of microhaplotypes (>250 reads/target) and drug resistance targets by parasite density bins. **C.** Standard error in haplotype proportion between monoclonal and polyclonal replicates. The number of replicates per sample and the complexity of infections are shown. **D.** Distribution of multiplicity of infection in field samples from Mozambique. **E.** Distribution of expected heterozygosity and the number of effective alleles in three provinces of southern Mozambique. **F.** Correlation of expected heterozygosity of microhaplotypes in the global population and the observed heterozygosity in the Mozambique samples. **G.** Visualization of microhaplotypes and SNP barcodes in polyclonal infections. Rows are samples and columns represent loci. For the microhaplotypes, each allele is represented by a unique colour at that locus and the size of the stacked bar is proportional to the within host proportion of the allele, generating a “molecular rainbow”.

## DISCUSSION

In this study, a bespoke bioinformatic pipeline was developed and validated to identify thousands of globally diverse, multiallelic microhaplotypes throughout the *P. falciparum* genome - the heterozygome. Using multiplex PCR, we were able to simultaneously sequence 100 heterozygome microhaplotypes and key drug resistance targets in a single reaction with consistent, deep coverage, detecting and quantifying minority alleles in DBS samples down to 10 parasites/µL of blood and outperforming whole genome sequences obtained from 50 times more total reads. Data from laboratory controls and in silico analyses indicate that this approach allows for better estimation of genetic diversity within and genetic relatedness between polyclonal samples when compared with panels of SNPs, making it a promising tool for studying the transmission dynamics of *P. falciparum*.

Measuring genetic relatedness between malaria infections is a fundamental step towards translating genomic data into operationally relevant information on transmission dynamics and tracking parasite flow [10,44,45]. The utility of genetic data in evaluating relatedness is largely driven by the diversity of the markers used, with greater diversity generally giving better resolution. When comparing individual parasites to each other, a sufficient number (e.g. a few hundred) of moderately diverse SNPs can theoretically provide sufficient resolution for comparisons, because combinations of many SNPs occurring within an individual parasite may be informative [44]. In practice, the ability of SNP panels to measure meaningful differences in relatedness is limited in many settings because a large proportion of infections are polyclonal, reducing the multiplicative benefit of numerous markers since phased combinations belonging to individual parasites are not directly observed. In such settings, the incremental value of increasing the diversity of each locus beyond that obtained from biallelic SNPs becomes greater and may be necessary to obtain meaningful information. Intuitively, this is because in the absence of phasing, comparisons between polyclonal infections are often reduced to determining whether or not alleles are shared at each locus; when all possible alleles at a locus are present (2 in the case of most SNPs) there will always be alleles shared with all other infections. As a result, many studies have only compared monoclonal infections, limiting power and potentially introducing bias into analyses. Multiallelic markers such as microsatellites have been used to overcome these limitations in the past but are cumbersome; microhaplotypes accomplish a similar objective using current sequencing technology by unambiguously phasing multiple SNPs that are close enough to be sequenced in a single read [16,17,46]. As demonstrated here, the combinations of SNPs within properly selected microhaplotypes can provide substantial diversity, dramatically outperforming independent SNPs in the ability to estimate relatedness between polyclonal infections. Furthermore, we have shown that deep sequencing of microhaplotypes provides sensitive detection and accurate quantification of minority alleles, potentially allowing for in silico phasing of microhaplotypes in mixed infections and allowing for even higher resolution comparisons.

While we show the application of this approach using ∼100 microhaplotypes as an example, the number and composition of markers can be tailored to specific questions of interest and target populations. For example, to understand within host diversity and genetic relatedness on a timescale that is relevant to directing and evaluating interventions (e.g. driven by recombination events occurring over months to a few years) high diversity microhaplotypes should provide the required resolution. Based on our analysis, many of these loci are diverse across multiple geographic settings. If the goal is to perform spatial assignment of infections, microhaplotypes with high genetic differentiation between infections from relevant geographic regions should be considered. These may be more specific to particular contexts, as signals of genetic differentiation are more varied geographically and at different spatial scales. Thus, the availability of globally diverse windows across different transmission zones should allow the development of a core panel, with the potential to add a subset of highly differentiated microhaplotypes for geographic regions of interest. It should be noted that many of the microhaplotypes identified herein may be under balancing or directional selection. As such, they may not be appropriate for analyses which mandate neutral markers [47]. If neutrality is preferred but not strictly required, the statistical power of having diverse markers must be balanced against this preference. Another consideration is that this study evaluated microhaplotypes that can be sequenced with 150bp paired end reads; improvements in long-read targeted sequencing may additionally allow the consideration of longer microhaplotypes or minihaplotypes (∼10kb) for increased genetic resolution and greater flexibility to incorporate complex and diverse regions of the genome [48].

This analysis has identified a set of generally informative markers and has begun to evaluate the utility of using microhaplotypes for high resolution genotyping of low-density and polyclonal infections. However, recognizing the full potential of genetic data generated from this approach will require additional improvements in downstream analysis. Few analytical tools directly address the challenges of incorporating multiallelic data or evaluation of polyclonal infections; even fewer encompass both. For example, to our knowledge there are no established methods for calculating basic statistics such as population level allele frequency or formally calculating genetic distance between polyclonal infections using multiallelic markers, forcing the use of ad hoc metrics such as those used here and in prior analyses [14]. Near term investment in developing these tools will be necessary to take full advantage of the rich data generated from *P. falciparum* microhaplotypes in translating genomic data into meaningful epidemiologic intelligence. The development of appropriate analytical frameworks will also facilitate the rational selection of informative microhaplotype panels, e.g. determining how many and which microhaplotypes provide adequate resolution to answer a set of questions in a range of epidemiologic settings. Availability of the appropriate amplicon panels and calibrated tools will bring us much closer to mapping epidemiologic and genetic data onto operationally relevant measures such as changes in transmission intensity, evaluation of the impact of interventions, accurate classification of local and imported infections, and evaluation of transmission chains [49,50].

Incorporating genomics into malaria surveillance in endemic settings will not be trivial, but the availability of efficient and information rich strategies for generating genomic data from routinely collected field samples, such as the one evaluated here, is an important step towards this goal. The ability to simultaneously obtain deep sequence data from targets informing drug resistance and multiple aspects of transmission epidemiology should allow for streamlining and standardization of laboratory and bioinformatic pipelines, allowing these technologies to be transferred more rapidly to endemic settings. Additional components not evaluated here, such as species identification, an expanded number of candidate and validated molecular markers of drug resistance, and markers of diagnostic resistance (e.g. *hrp2/3* deletions) would enable a larger set of use cases to be addressed simultaneously. With such a panel and appropriate analytical tools, an integrated approach to genomic surveillance allowing for routine generation of these valuable data in the areas where they are needed most may be closer to becoming a reality.

## FUNDING

This work was supported by the Bill and Melinda Gates Foundation (Award Number OPP1132226) and by the Chan Zuckerberg Biohub. BG is a Chan Zuckerberg Biohub investigator.

## ACKNOWLEDGMENTS

The authors would like to thank the participants in the Mozambique studies, the participants’ parents and guardians and the field study team who performed sample and data collection. This publication uses data from the MalariaGEN *P. falciparum* Community Project (www.malariagen.net/projects/p-falciparum-community-project). MalariaGEN’s genome sequencing was performed by the Wellcome Trust Sanger Institute and the Community Projects is coordinated by the MalariaGEN Resource Centre with funding from the Wellcome Trust (098051, 090770).

## CONFLICTS OF INTEREST

The authors declare no competing interests. The funders of the study had no role in the study design, data collection, data interpretation, or writing of the manuscript. The authors had final responsibility for the decision to submit for publication.

## Additional files

**Supplementary Method:** laboratory protocol and amplicon analysis

**Supplementary Figure 1.** Strain composition of mock mixture controls

**Supplementary Figure 2.** Coverage of whole genome sequencing of mixture control samples.

**Supplementary figure 3.** Coverage and sensitivity of microhaplotype sequences extracted from WGS of mixture control samples.

**Supplementary Table 1**. List of publicly available whole genome sequences used in this study

**Supplementary Table 2**. The heterozygome of *P. falciparum*

**Supplementary Table 2**. Description of *P. falciparum* microhaplotypes

## Supplementary Methods: laboratory protocol and amplicon analysis

### 1. Primer pool

Primers were designed for 100 selected genomic regions of the microhaplotypes and drug resistance markers using the CleanPlex^®^ algorithm and synthesized by Paragon Genomics Inc, USA (Table A). A list of targets and primers is included in Supplementary Table 3.

**Table A.** Description of primers.

| *panel* | *Organism* | *Number of pools* | *Primer conc.* | *Number of targets* | *Average length* | *Library length* |
| --- | --- | --- | --- | --- | --- | --- |
| PGD268 | Pf | 1 | 5X | 100 | 228bp | 300-400bp |

### 2. Primary multiplexed PCR amplification

Amplification of the 200-plex oligo pool was performed with some modifications of the CleanPlex® protocol (Paragon Genomics Inc, USA). Two sets of PCR conditions were optimized for low (i.e. ≤ 100 parasites per µL blood, condition “C1”) and high (i.e. > 100 parasites per µL blood, condition “C5”) parasite density dried blood spot samples. A two-step primary multiplex PCR was carried out in a 10 µL reaction containing 6 µL genomic DNA (Table B and C). Primary PCR product was diluted with 10 μl of 1X TE buffer and SPRI (Solid Phase Reversible Immobilization) bead purified using a sample:bead volume ratio of 1:1.3 and resuspended in 10μL 1X TE buffer after the final wash. Following bead purification, 10 μL of CleanPlex^®^ digest master mix was added and incubated for 10 min at 37°C and followed by SPRI bead purification using a sample:bead volume ratio of 1:1.3 and resuspended in 10μL (Table D).

**Table B.** Master mix composition for the two PCR conditions.

| Master mix for the multiple PCR |  |  |  |
| --- | --- | --- | --- |
| C1 - condition |  | C5 - condition |  |
| Reagents* | Volume | Reagents* | Volume |
| 5X mPCR Mix | 2$\mu\text{L}$ | 5X mPCR Mix | 2$\mu\text{L}$ |
| 5X mPCR Primer | 2$\mu\text{L}$ | 5X mPCR Primer (1in 4 diluted) | 2$\mu\text{L}$ |
| DNA Sample** | 6$\mu\text{L}$ | DNA Sample** | 6$\mu\text{L}$ |
| total | 10$\mu\text{L}$ | total | 10$\mu\text{L}$ |

\*All reagents were supplied by Paragon Genomics Inc, CA, USA
\*\*DNA was extracted from dried blood spot samples using Tween-Chelex extraction protocol.

**Table C.** Cycling conditions.

| Thermal Cycling Conditions |  |  |  |  |  |  |  |  |  |
| --- | --- | --- | --- | --- | --- | --- | --- | --- | --- |
|  | C1 - condition |  |  |  |  | C5 - condition |  |  |  |
| Step | Temp | Time | Ramping | Cycles |  | Temp | Time | Ramping | Cycles |
| Initial Denaturation | 95°C | 10 min | - | 1 |  | 95°C | 10 min | - | 1 |
| Denaturation | 98°C | 15 sec | 3°C/s | 10 cycles |  | 98°C | 15 sec | 3°C/s | 30 cycles |
| Annealing/ Extension | 60°C | 5 min | 2°C/s |  |  | 60°C | 5 min | 2°C/s |  |
| Hold | 10°C | ∞ |  |  |  | 10 °C | ∞ |  |  |

**Table D.** Digestion reaction master mix.

| Reagents* | Volume |
| --- | --- |
| Nuclease-Free Water | 6µl |
| CP Reagent Buffer | 2µl |
| CP Digestion Reagent | 2µl |
| total | 10µl |

\*All reagents were supplied by Paragon Genomics Inc, CA, USA

### 3. Indexing PCR and pooling

A secondary indexing PCR was performed in a 40 µL reaction containing 10 µL of bead purified digested product, 18μL of nuclease-free water, 8μL of 5X secondary PCR master mix, and 5 µL of 10 µM TruSeq i5/i7 barcode primers. PCR was carried out using the following cycling conditions: 10 min at 95°C, 13 cycles for high density samples (or 15 cycles for low density samples) of 15 sec at 98°C and 75 sec at 60°C. Samples were SPRI bead purified and quantified by capillary electrophoresis. All samples were proportionally pooled based on the estimated concentration of the capillary electrophoresis. The final library was bead purified, assessed for quality on the Bioanalzyer (Agilent technologies, Santa Clara, CA) and sequenced with 150bp paired end clusters on the Illumina NextSeq 550 instrument (Illumina, San Diego, CA, USA).

### 4. Targeted amplicon analysis

The targeted amplicon data were analyzed using SeekDeep (v.2.6.6) [1]. Briefly, Illumina pair-end sample files were demultiplexed on target primers and paired end reads stitched into a single read while filtering for per base quality and expected target lengths to create a FASTQ file for each target-sample pair. The SeekDeep qluster step was used to create haplotypes per sample per target by comparing sequences and collapsing on low per base quality errors and low frequency error to help eliminate sequencing and low-level PCR error. Each haplotype was given a within sample within target frequency based on the number of reads clustered together and haplotypes marked as possibly chimeric if there were two possible parent sequences at a frequency of at least twice as much as the possible chimeric haplotype. SeekDeep processClusters step then handled the final processing of the haplotypes by removing the marked possibly chimeric haplotypes. The final haplotypes for each target were then compared across samples to get population frequency calculations and to generate final population haplotypes. To help further remove possible artifact that was either introduced by PCR or by cross contamination between samples in either library prep or within the Illumina machine [2] several additional post processing filtering steps were performed during the SeekDeep processClusters step. Haplotypes that were either one SNP or one indel different from a within sample haplotype that was found with at least 10 times greater frequency were removed unless that haplotype was seen in another sample as a major haplotype. Haplotypes that only appeared in one sample and that were one difference off another haplotype within sample were also removed to reduce the detection of PCR jackpot events that happen early in PCR. Haplotypes were also removed if below a 2% frequency.

---

**Supplementary Figure 1.**
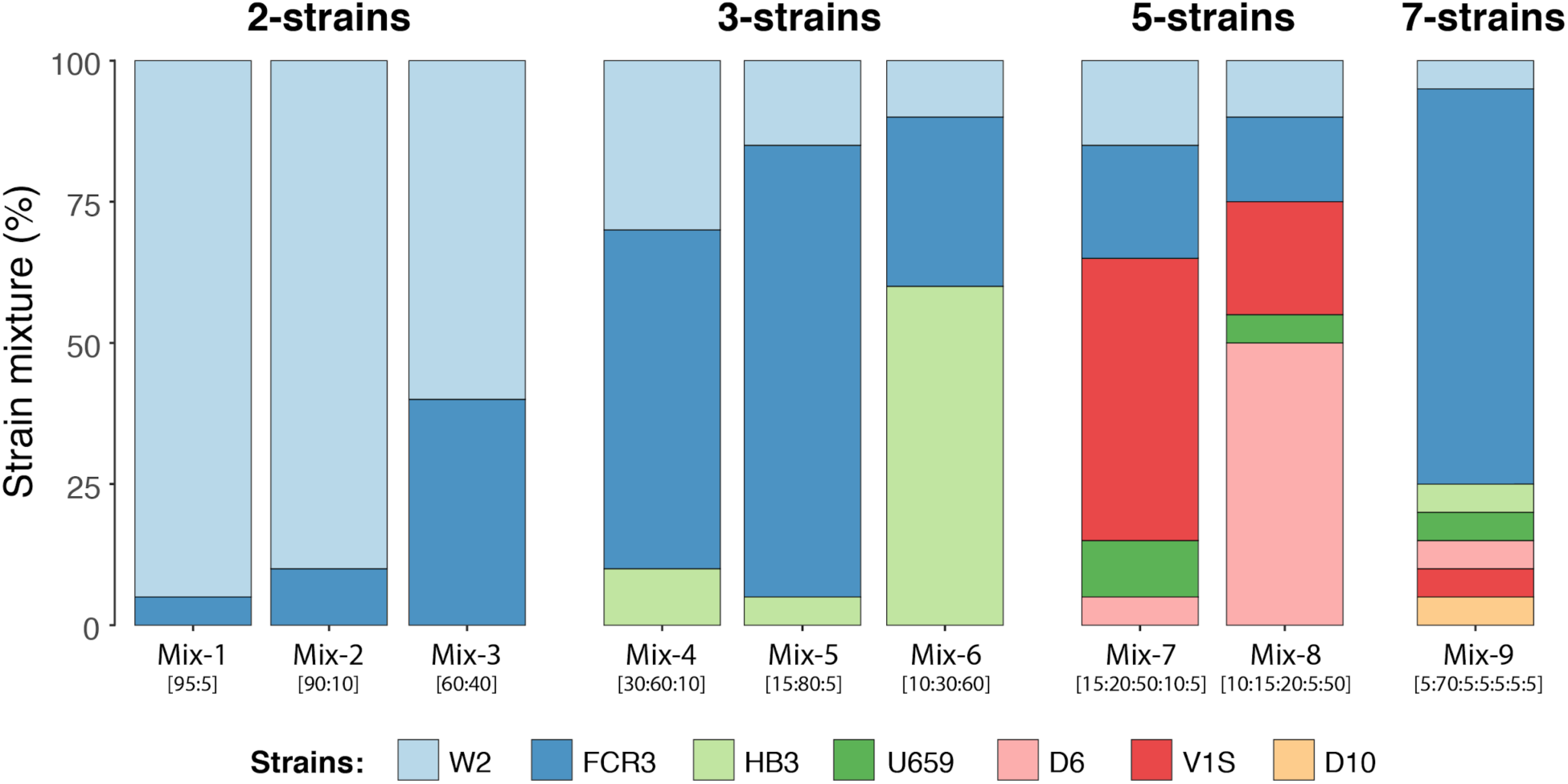
Strain composition of mock mixture controls. Different numbers of strains were mixed at different proportions to simulate polyclonal infections.

**Supplementary Figure 2.**
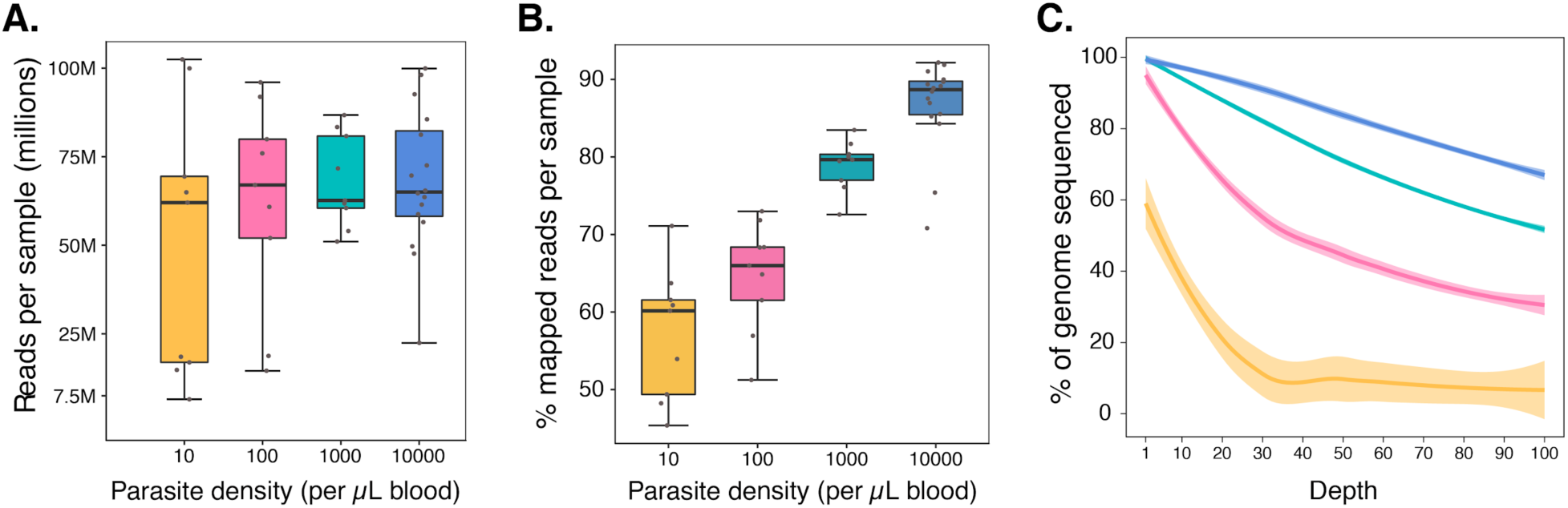
Coverage of whole genome sequencing of mixture control samples. **A.** Box plot showing the total number of reads sequenced by parasite density. **B.** Boxplot showing the proportion of reads mapped to the core *P. falciparum* genome. **C.** The percentage of the core *P. falciparum* genome covered by a minimum read depth is shown; colors correspond to parasite density in panels A and B.

**Supplementary figure 3.**
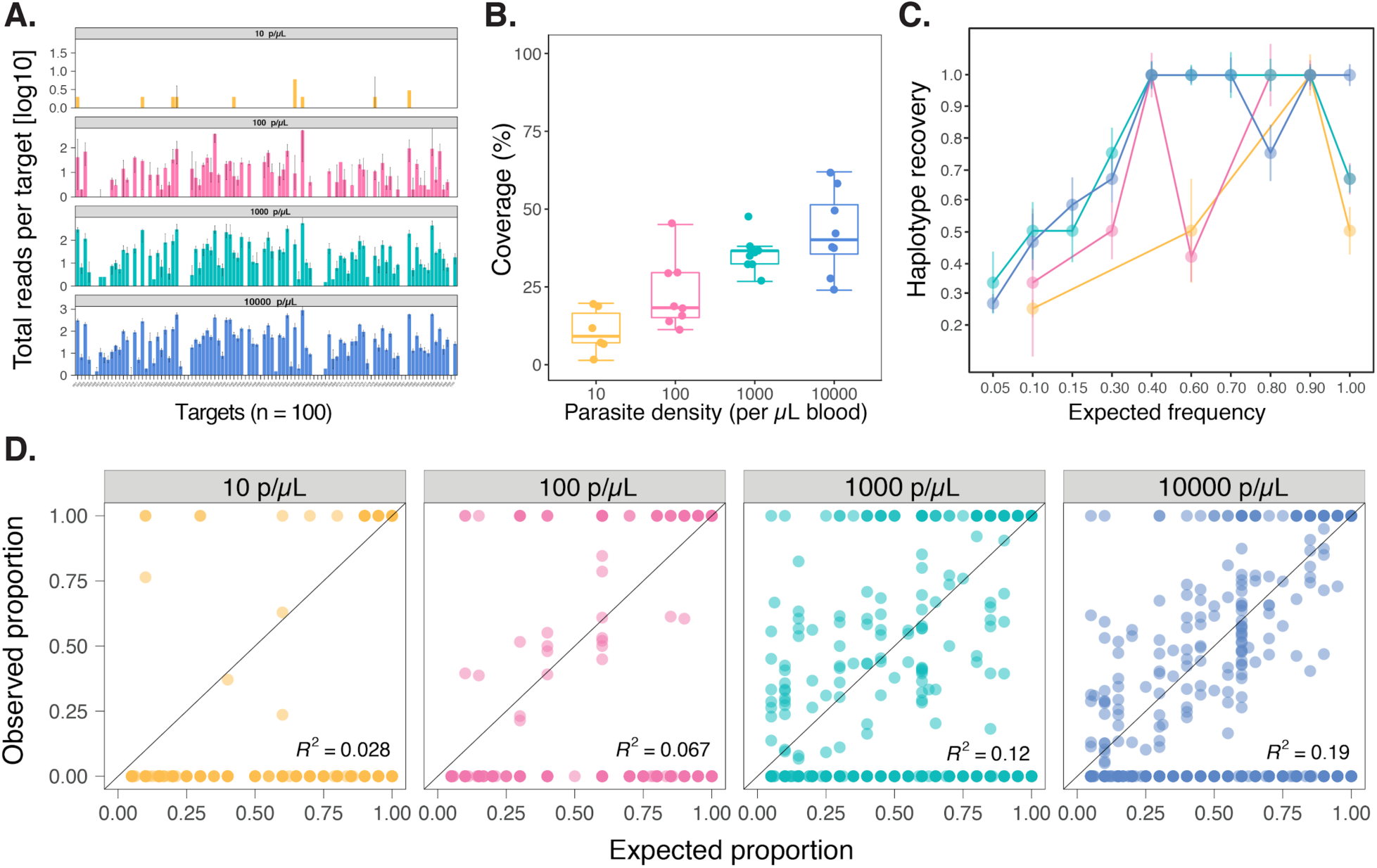
Coverage and sensitivity of microhaplotype sequences extracted from WGS of mixture control samples. **A.** Average number of fully spanning reads per microhaplotype. The median (bars) and interquartile range (error bars) are shown. **B.** Boxplot summarising the coverage of full-length microhaplotype loci extracted from WGS data is shown by parasite density. **C.** Microhaplotype recovery in WGS extracted reads shown by parasite density. **D.** Correlation of expected and observed frequencies of microhaplotypes by parasite density is shown.

## REFERENCES

1. Hamilton WL, Amato R, van der Pluijm RW, Jacob CG, Quang HH, Thuy-Nhien NT, et al. Evolution and expansion of multidrug-resistant malaria in southeast Asia: a genomic epidemiology study. Lancet Infect Dis. 2019;19:943–51.

2. Neafsey DE, Volkman SK. Malaria Genomics in the Era of Eradication. Cold Spring Harb Perspect Med [Internet]. 2017;7. Available from: http://dx.doi.org/10.1101/cshperspect.a025544

3. Kwiatkowski D. Malaria genomics: tracking a diverse and evolving parasite population. Int Health. 2015;7:82–4.

4. Viriyakosol S, Siripoon N, Petcharapirat C, Petcharapirat P, Jarra W, Thaithong S, et al. Genotyping of Plasmodium falciparum isolates by the polymerase chain reaction and potential uses in epidemiological studies. Bull World Health Organ. 1995;73:85–95.

5. Anderson TJ, Su XZ, Bockarie M, Lagog M, Day KP. Twelve microsatellite markers for characterization of Plasmodium falciparum from finger-prick blood samples. Parasitology. 1999;119 (Pt 2):113–25.

6. Anderson TJ, Haubold B, Williams JT, Estrada-Franco JG, Richardson L, Mollinedo R, et al. Microsatellite markers reveal a spectrum of population structures in the malaria parasite Plasmodium falciparum. Mol Biol Evol. 2000;17:1467–82.

7. Daniels R, Volkman SK, Milner DA, Mahesh N, Neafsey DE, Park DJ, et al. A general SNP-based molecular barcode for Plasmodium falciparum identification and tracking. Malar J. 2008;7:223.

8. Nkhoma SC, Nair S, Al-Saai S, Ashley E, McGready R, Phyo AP, et al. Population genetic correlates of declining transmission in a human pathogen. Mol Ecol. 2013;22:273–85.

9. Chang H-H, Wesolowski A, Sinha I, Jacob CG, Mahmud A, Uddin D, et al. Mapping imported malaria in Bangladesh using parasite genetic and human mobility data. Elife [Internet]. 2019;8. Available from: http://dx.doi.org/10.7554/eLife.43481

10. Wesolowski A, Taylor AR, Chang H-H, Verity R, Tessema S, Bailey JA, et al. Mapping malaria by combining parasite genomic and epidemiologic data. BMC Med. 2018;16:190.

11. Apinjoh TO, Ouattara A, Titanji VPK, Djimde A, Amambua-Ngwa A. Genetic diversity and drug resistance surveillance of Plasmodium falciparum for malaria elimination: is there an ideal tool for resource-limited sub-Saharan Africa? Malar J. 2019;18:217.

12. Figan CE, Sá JM, Mu J, Melendez-Muniz VA, Liu CH, Wellems TE. A set of microsatellite markers to differentiate Plasmodium falciparum progeny of four genetic crosses. Malar J. 2018;17:60.

13. Daniels RF, Schaffner SF, Wenger EA, Proctor JL, Chang H-H, Wong W, et al. Modeling malaria genomics reveals transmission decline and rebound in Senegal. Proc Natl Acad Sci U S A. 2015;112:7067–72.

14. Tessema S, Wesolowski A, Chen A, Murphy M, Wilheim J, Mupiri A-R, et al. Using parasite genetic and human mobility data to infer local and cross-border malaria connectivity in Southern Africa. Elife [Internet]. 2019;8. Available from: http://dx.doi.org/10.7554/eLife.43510

15. Roh ME, Tessema SK, Murphy M, Nhlabathi N, Mkhonta N, Vilakati S, et al. High Genetic Diversity of Plasmodium falciparum in the Low-Transmission Setting of the Kingdom of Eswatini [Internet]. The Journal of Infectious Diseases. 2019. p. 1346–54. Available from: http://dx.doi.org/10.1093/infdis/jiz305

16. Kidd KK, Pakstis AJ, Speed WC, Lagace R, Chang J, Wootton S, et al. Microhaplotype loci are a powerful new type of forensic marker. Forensic Science International: Genetics Supplement Series. 2013;4:e123–4.

17. Oldoni F, Kidd KK, Podini D. Microhaplotypes in forensic genetics. Forensic Sci Int Genet. 2019;38:54–69.

18. Nag S, Dalgaard MD, Kofoed P-E, Ursing J, Crespo M, Andersen LO, et al. High throughput resistance profiling of Plasmodium falciparum infections based on custom dual indexing and Illumina next generation sequencing-technology. Sci Rep. 2017;7:2398.

19. Talundzic E, Ravishankar S, Kelley J, Patel D, Plucinski M, Schmedes S, et al. Next-Generation Sequencing and Bioinformatics Protocol for Malaria Drug Resistance Marker Surveillance. Antimicrob Agents Chemother [Internet]. 2018;62. Available from: http://dx.doi.org/10.1128/AAC.02474-17

20. Lerch A, Koepfli C, Hofmann NE, Messerli C, Wilcox S, Kattenberg JH, et al. Development of amplicon deep sequencing markers and data analysis pipeline for genotyping multi-clonal malaria infections. BMC Genomics. 2017;18:864.

21. Neafsey DE, Juraska M, Bedford T, Benkeser D, Valim C, Griggs A, et al. Genetic Diversity and Protective Efficacy of the RTS,S/AS01 Malaria Vaccine [Internet]. New England Journal of Medicine. 2015. p. 2025–37. Available from: http://dx.doi.org/10.1056/nejmoa1505819

22. Juliano JJ, Porter K, Mwapasa V, Sem R, Rogers WO, Ariey F, et al. Exposing malaria in-host diversity and estimating population diversity by capture-recapture using massively parallel pyrosequencing. Proc Natl Acad Sci U S A. 2010;107:20138–43.

23. Miller RH, Hathaway NJ, Kharabora O, Mwandagalirwa K, Tshefu A, Meshnick SR, et al. A deep sequencing approach to estimate Plasmodium falciparum complexity of infection (COI) and explore apical membrane antigen 1 diversity [Internet]. Malaria Journal. 2017. Available from: http://dx.doi.org/10.1186/s12936-017-2137-9

24. Gandhi K, Thera MA, Coulibaly D, Traoré K, Guindo AB, Doumbo OK, et al. Next generation sequencing to detect variation in the Plasmodium falciparum circumsporozoite protein. Am J Trop Med Hyg. 2012;86:775–81.

25. Early AM, Daniels RF, Farrell TM, Grimsby J, Volkman SK, Wirth DF, et al. Detection of low-density Plasmodium falciparum infections using amplicon deep sequencing. Malar J. 2019;18:219.

26. Verity R, Aydemir O, Brazeau NF, Watson OJ, Hathaway NJ, Mwandagalirwa MK, et al. The Impact of Antimalarial Resistance on the Genetic Structure of Plasmodium falciparum in the DRC [Internet]. bioRxiv. 2019 [cited 2019 Nov 15]. p. 656561. Available from: https://www.biorxiv.org/content/10.1101/656561v1.abstract

27. Aydemir O, Janko M, Hathaway NJ, Verity R, Mwandagalirwa MK, Tshefu AK, et al. Drug-Resistance and Population Structure of Plasmodium falciparum Across the Democratic Republic of Congo Using High-Throughput Molecular Inversion Probes. J Infect Dis. 2018;218:946–55.

28. Pearson RD, Amato R, Kwiatkowski DP. An open dataset of Plasmodium falciparum genome variation in 7,000 worldwide samples. bioRxiv [Internet]. biorxiv.org; 2019; Available from: https://www.biorxiv.org/content/10.1101/824730v1.abstract

29. Amambua-Ngwa A, Amenga-Etego L, Kamau E, Amato R, Ghansah A, Golassa L, et al. Major subpopulations of Plasmodium falciparum in sub-Saharan Africa. Science. 2019;365:813–6.

30. Benson G. Tandem repeats finder: a program to analyze DNA sequences. Nucleic Acids Res. 1999;27:573–80.

31. Otto TD, Böhme U, Sanders M, Reid A, Bruske EI, Duffy CW, et al. Long read assemblies of geographically dispersed Plasmodium falciparum isolates reveal highly structured subtelomeres. Wellcome Open Res. 2018;3:52.

32. Harris RS. Improved pairwise alignment of genomic DNA [Internet]. The Pennsylvania State University; 2007. Available from: http://www.bx.psu.edu/~rsharris/rsharris_phd_thesis_2007.pdf

33. Teyssier NB, Chen A, Duarte EM, Sit R, Greenhouse B, Tessema SK. Optimization of whole-genome sequencing of Plasmodium falciparum from low-density dried blood spot samples [Internet]. bioRxiv. 2019 [cited 2019 Nov 13]. p. 835389. Available from: https://www.biorxiv.org/content/10.1101/835389v1.abstract

34. Hathaway NJ, Parobek CM, Juliano JJ, Bailey JA. SeekDeep: single-base resolution de novo clustering for amplicon deep sequencing. Nucleic Acids Res. 2018;46:e21.

35. Li H. Aligning sequence reads, clone sequences and assembly contigs with BWA-MEM. arXiv preprint 1303 3997 [Internet]. 2013; Available from: http://arxiv.org/abs/1303.3997

36. Auwera GA, Carneiro MO, Hartl C, Poplin R, del Angel G, Levy-Moonshine A, et al. others.(2013). From FastQ data to high-confidence variant calls: the genome analysis toolkit best practices pipeline. Curr Protoc Bioinformatics. :11–10.

37. Dara A, Drábek EF, Travassos MA, Moser KA, Delcher AL, Su Q, et al. New var reconstruction algorithm exposes high var sequence diversity in a single geographic location in Mali. Genome Med. 2017;9:30.

38. Parobek CM, Parr JB, Brazeau NF, Lon C, Chaorattanakawee S, Gosi P, et al. Partner-Drug Resistance and Population Substructuring of Artemisinin-Resistant Plasmodium falciparum in Cambodia. Genome Biol Evol. Oxford University Press; 2017;9:1673–86.

39. Cerqueira GC, Cheeseman IH, Schaffner SF, Nair S, McDew-White M, Phyo AP, et al. Longitudinal genomic surveillance of Plasmodium falciparum malaria parasites reveals complex genomic architecture of emerging artemisinin resistance. Genome Biol. 2017;18:78.

40. Baniecki ML, Faust AL, Schaffner SF, Park DJ, Galinsky K, Daniels RF, et al. Development of a single nucleotide polymorphism barcode to genotype Plasmodium vivax infections. PLoS Negl Trop Dis. 2015;9:e0003539.

41. Kumar S, Mudeppa DG, Sharma A, Mascarenhas A, Dash R, Pereira L, et al. Distinct genomic architecture of Plasmodium falciparum populations from South Asia. Mol Biochem Parasitol. 2016;210:1–4.

42. Krijthe JH. Rtsne: T-distributed stochastic neighbor embedding using Barnes-Hut implementation. R package version 0 13, URL https://githubcom/jkrijthe/Rtsne. 2015;

43. Chang H-H, Worby CJ, Yeka A, Nankabirwa J, Kamya MR, Staedke SG, et al. THE REAL McCOIL: A method for the concurrent estimation of the complexity of infection and SNP allele frequency for malaria parasites. PLoS Comput Biol. 2017;13:e1005348.

44. Taylor AR, Jacob PE, Neafsey DE, Buckee CO. Estimating Relatedness Between Malaria Parasites. Genetics. 2019;212:1337–51.

45. Tessema SK, Raman J, Duffy CW, Ishengoma DS, Amambua-Ngwa A, Greenhouse B. Applying next-generation sequencing to track falciparum malaria in sub-Saharan Africa. Malar J. 2019;18:268.

46. Baetscher DS, Clemento AJ, Ng TC, Anderson EC, Garza JC. Microhaplotypes provide increased power from short-read DNA sequences for relationship inference. Mol Ecol Resour. 2018;18:296–305.

47. Volkman SK, Neafsey DE, Schaffner SF, Park DJ, Wirth DF. Harnessing genomics and genome biology to understand malaria biology. Nat Rev Genet. 2012;13:315–28.

48. Mantere T, Kersten S, Hoischen A. Long-Read Sequencing Emerging in Medical Genetics. Front Genet. 2019;10:426.

49. WHO. Technical consultation on the role of parasite and anopheline genetics in malaria surveillance. Malaria Policy Advisory Committee Meeting [Internet]. 2019; Available from: https://www.who.int/malaria/mpac/mpac-october2019-session7-report-consultation-on-genomics.pdf

50. Dalmat R, Naughton B, Kwan-Gett TS, Slyker J, Stuckey EM. Use cases for genetic epidemiology in malaria elimination. Malar J. 2019;18:163.

## References

1. Hathaway NJ, Parobek CM, Juliano JJ, Bailey JA. SeekDeep: single-base resolution de novo clustering for amplicon deep sequencing. Nucleic Acids Res. 2018; 46(4):e21.

2. Costello M, Fleharty M, Abreu J, et al. Characterization and remediation of sample index swaps by non-redundant dual indexing on massively parallel sequencing platforms. BMC Genomics. 2018; 19(1):332.

